# Molecular dynamics simulations reveal DNA gate opening mechanisms for *M. smegmatis* topoisomerase 1A

**DOI:** 10.1101/2025.02.13.638100

**Authors:** Deepesh Sigdel, Maria Mills

## Abstract

Type 1A topoisomerases relax torsional strain in DNA via a strand passage mechanism in which a protein mediated DNA gate must open during the enzyme’s catalytic cycle. This gate-open conformational state of the enzyme has been challenging to observe via experimental methods. In this study, we first used equilibrium Molecular Dynamics simulations to probe the structural properties of the gate closed state for the DNA-free apo system and a system with a ssDNA bound at the DNA binding site. For both systems, we followed the equilibrium simulations with Umbrella Sampling simulations. Umbrella sampling allowed us to bias the protein to adopt a gate-open state to study the properties of this conformation, as well as the pathways leading to it. We observed that several electrostatic interactions contribute to the closed-state stability of the protein which were broken during the gate opening. The gate opening comprised of three major domain motions that were determined from simulation trajectories and Principal Component Analysis. Finally, umbrella sampling results combined with the Weighted Histogram Analysis Method allowed us to reconstruct the free energy profiles of gate opening for all simulations.

**SIGNIFICANCE:** Multi-drug resistant bacterial strains necessitate newer ways of targeting bacteria. Type 1A topoisomerases are potential anti-bacterial drug targets. Current strategies are to either block the protein binding to DNA or prevent the cut DNA being religated during the catalytic cycle. A possible alternative to these methods would be to trap the protein at an open conformation. Structural information about the transient open state is crucial to this approach. Results of this study provide mechanistic details of gate opening. This information furthers the understanding of the topoisomerase catalytic cycle which will aid in anti-topoisomerase drug design efforts.

## INTRODUCTION

DNA topoisomerases are important enzymes that resolve topological issues in DNA including knots, precatenanes, excess supercoils and similar. These DNA structures occur as a result of various cellular processes such as DNA replication and transcription [1]. Topoisomerases are classified into two major families, Type 1 and Type 2 depending on whether they cleave one or both strands of DNA for their catalytic activity[2]. Type 1A topoisomerases are members of the type 1 topoisomerases, and use a strand passage mechanism in which an intact ssDNA strand – commonly referred to as the T-strand - is passed through a cut in the other strand, called the G-strand. The protein must undergo a large conformational change that moves the cut ends of the G-strand apart to create an opening for the T-strand to pass through.

Type 1A topoisomerases share a conserved N-terminal structure formed by domains 1 - 4 (D1-D4) that exhibit a toroidal padlock type structure (Figure 1(A)). The C-terminal domains (grey, Figure 1(A,B)) however vary between different type 1A topoisomerases[3]. During protein activity, the G-strand binds across D1 and D4 domains while a catalytic Tyrosine from D3 cleaves the ssDNA forming a 5^′^ phosphotyrosyl linkage with the cut end. To pass a T-strand through this cut, a conformational change presumably occurs where D3 moves away from D4. Based on the proposed protein activity models, in a gate open state, D3 moves away from both D1 and D4[4]. This conformational change does not require ATP [5].

**Figure 1:**
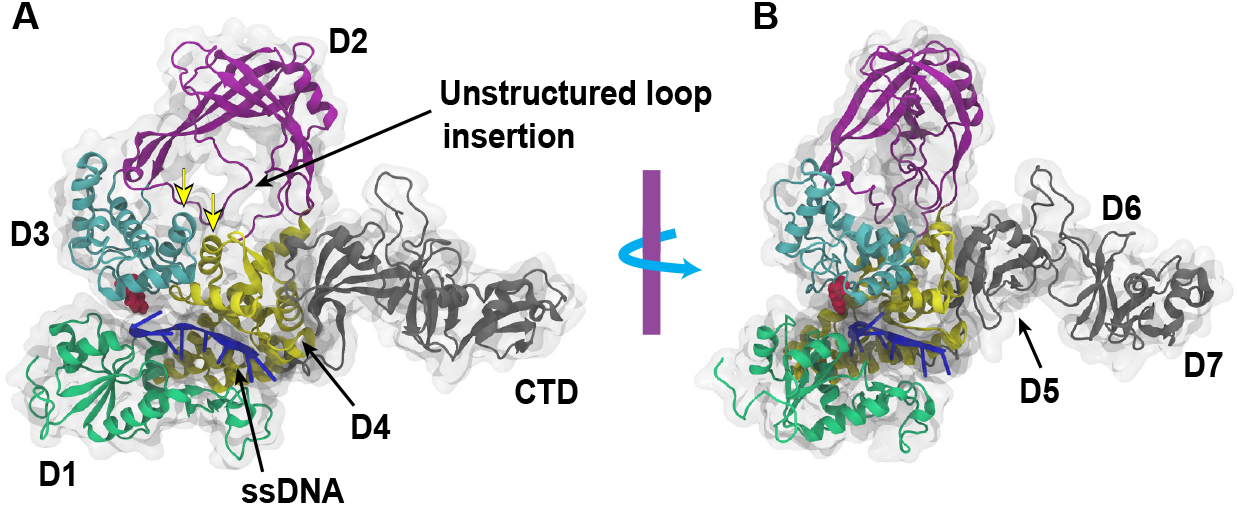
Msm Top1A structure and features. (A) Msm Top1A structure adapted from PDB ID: 6PCM[15]. N-terminal toroid domains colored D1-green, D2-purple, D3-cyan, D4-yellow. C-terminal domains collectively colored grey. This face is referred to as the front face of the protein throughout this work. An unstructured loop is seen protruding out of D2. The loop extends across the interface alpha helices (cyan and yellow helices pointed by yellow arrows) and into the toroid cavity. (B) A slightly rotated view of the same structure showing how the C-terminal tail domains extend away from the plane (or face) of the protein. Domain D8, and an unstructured tail were not present in this structure. A boundary interface exists between D3, D1 and D4 domains. This interface is referred to as the gate in this study. Catalytic Tyrosine (red) is seen at the interface protruding towards the ssDNA in both panels.

The type 1A topoisomerase mechanism of activity has been well studied using several biochemical [6], [7] and biophysical techniques [8]–[10]. Single molecule experiments have shown that these proteins relax excessive supercoils via a strand passage mechanism in steps of one excess turn at a time[8]. Other studies have shown that the proteins must undergo significant conformational changes during the reaction cycle[11], [12]. There have been many structural studies throughout the years looking at type 1A topoisomerases [13]–[15]. Structures of intermediate states had been elusive until a recent structural study on Human Top 3*β*[16] revealed an open conformation. This elusive nature of the open state suggests that this state is highly transient and less stable than the closed-state. Single molecule experiments have also contributed to the understanding of the protein mechanisms regarding the gate open state. It has been demonstrated that the application of force can bias the protein to dwell longer at these transient gate-open states[17]. Individual protein-mediated DNA gate openings were directly observed using single molecule magnetic, and optical tweezers [17], [18]. These experiments demonstrated that the protein domains undergo large conformational changes upon DNA cleavage with DNA gate opening >5 nm [17], [18] under application of pN forces.

Little is known about type 1A topoisomerase structures at different stages of the reaction cycle and the mechanisms by which they occur. The transient nature of the open state makes visualizing the conformational dynamics challenging. Therefore, a dynamic study is essential to be able to capture such processes. Molecular dynamics (MD) simulations is one way to study such mechanisms. MD simulations are regularly employed to probe protein conformational changes, domain reorganization, folding-unfolding pathway and much more[19]. MD simulations can incorporate different biasing techniques to force a protein to explore alternative conformational states as well.

In this work, we used MD simulations to probe the dynamic features of the *M. smegmatis* type 1A topoisomerase (Msm Top1A) protein structure. Msm Top1A is closely related to *Mycobacterium tuberculosis* Top1A - a validated anti-topoisomerase drug target[20], [21]. A better understanding of the Msm Top1A structural dynamics will aid in such drug discovery efforts. Using equilibrium simulations, we identified stable and mobile regions of the protein as well as residues important for the closed state stability. Using Umbrella Sampling (US) simulations, we explored the structural changes that occur during gate opening. We computed one and two-dimensional free energies of gate opening as well as identified important domain motions during this process. Principal component analysis results support the domain motions as being the major functional motions of the gate opening. Taken together, the findings of this study shed light on the energetics and dynamics of the elusive protein-mediated DNA gate opening.

## METHODS

Starting structures for all systems were based on Msm Top1A structure from the Protein Data Bank (PDB ID: 6PCM[15]). The protein crystallographic structure as solved lacks the last domain D8, and an unstructured tail. This slightly truncated structure was used in this study as we don’t expect these missing domains to influence gate opening. Missing residues within the truncated protein structure were modeled using Modeller via ChimeraX[22]. Among the Modeller generated structures, the structure with the lowest normalized Discrete Optimized Protein Energy Score(z-DOPE) was selected as the starting structure to build simulation systems. Systems were prepared using VMD. Apo protein and protein with ssDNA at the active site binding pocket were both solvated with TIP3P water molecules in a water box with at least 10 *Å* padding in all directions. Charges were neutralized and ion concentration set to 150 mM NaCl. The total atom count for each system was ∼ 186k and 206k respectively. For simulations with ssDNA at the active site pocket (+ssDNA), the longer ssDNA strand from the crystal structure was trimmed leaving only 8 nucleotides with sequence 5^′^ − *TTCCGCTT* − 3^′^ in place.

All MD simulations were atomistic in scale and run using NAMD 2.14 or NAMD 3.0 [23], [24]. Systems were energy minimized for at least 10000 steps using a conjugate gradient algorithm. Energy minimization was followed by isothermal-isobaric (NPT) equilibration at 1 bar and 310*K* using Langevin dynamics. Equilibration in the canonical ensemble (NVT) followed NPT equilibrations. Interactions were governed by CHARMM36[25] force field. Non-bonded interactions were truncated smoothly at 12 *Å* with a switching function activating at 10 *Å*. All electrostatics were treated by Particle Mesh Ewald (PME) method. For both systems, the total equilibration simulation time was as much as 0.2 *µs*. Runs were terminated once the Root Mean Square Deviation (RMSD) values for the core domains of the protein (D1-D4) had stabilized (ΔRMSD < 2*Å*). Both RMSD and time averaged RMSD per residue (Root Mean Square Fluctuation, RMSF) calculations were done using Python MDAnalysis libraries [26], [27]. Two further conformations were generated for each system where the unstructured loop was moved out of the cavity. These systems are referred to as apo_1 and +ssDNA_1 respectively.

PyMol was used to generate the electrostatic potential map of the protein as well as the figure showing the protein surface painted with the respective colors in Figure 3(A-D). The total nonbonded interaction energies between residues were calculated using VMD NAMDEnergy plugin with ϵ*_r_* = 70. The approximate D3 swiveling angle is calculated as the angle between *C*_*α*_ residues of Y339 (the active site Tyrosine), S516 and E523 (D4 interface helix residues). The latter two residues are found on the interface alpha-helix on the opposite side of the gate from the active site Tyrosine. A unit vector through S516 to E523 can be represented by 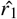, and from S516 to Y339 can be written as 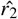. The angle between the two unit vectors is then calculated as cos 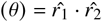 for all frames of the MD trajectories. The helix containing S516 and E523 (D4 *α*3) does not lose its structure throughout the MD simulation and is one of the most stable regions on the protein.

For gate opening, simulations were run using the biasing technique Umbrella Sampling(US)[28], [29]. For all runs, restraining potentials of the form 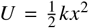 were used with spring constant *k* = 5 *k cal*/(*mol* · *Å*^2^). All US runs used a single reaction coordinate as shown in Figure 6(A). The entire reaction coordinate was split into 40 1*Å* windows which were used to steer the system from a closed to an open conformation. Reaction coordinate space was sampled every 2 *ps*. Each window was sampled for at least 4 *ns*. Simulations were ran for as much as 0.4 *µs*. For Free Energy calculations using the Weighted Histogram Analysis Method (WHAM)[30], last 90% of data from each window was retained. 2-dimensional WHAM calculations were carried out using the same code [30] as well.

All Principal Component Analysis (PCA) for the N-terminus *C*_*α*_ residues were done using ProDy libraries [31]–[33]. 20 lowest-energy uncorrelated modes were generated from each US run in this manner. VMD’s Normal Mode Wizard plugin was used to visualize all the modes. All trajectory snapshots except the electrostatics figure were generated using VMD[34] as well. NCBI Blast was used for multiple sequence alignment [35] and visualized using Jalview 2.11.4.1 [36] All plots were generated using Matplotlib [37].

## RESULTS

### Flexibility and mobility of the protein domains during equilibrium simulations

To determine the structural dynamics of Msm Top1A, we first ran equilibrium atomistic MD simulations on DNA-free (apo) Msm Top1A structure and the system with ssDNA at the binding site (+ssDNA). For all unrestrained equilibrium runs (200 ns for apo and 140 ns for +ssDNA), the protein Root Mean Square Deviation (RMSD) was calculated as mentioned in Methods. RMSD plots in Figure 2(A) demonstrate higher stability of N-terminal domains in both systems compared to the mobile C-terminal domains.

**Figure 2:**
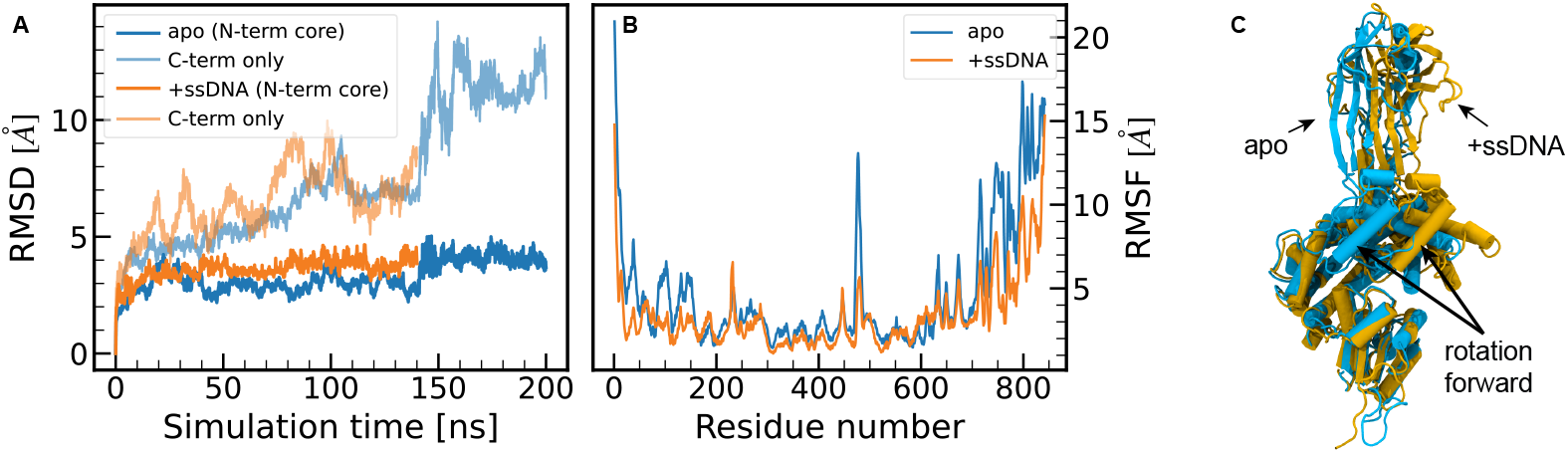
Analysis of equilibrium MD simulation trajectories for both systems. (A) RMSD trends over the total simulation time for core N-terminal domains (opaque lines), and C-terminal domains (translucent lines, same color) are shown for both apo, and +ssDNA systems. The high mobility of C-terminal residues can be seen as the high RMSD values that does not plateau. (B) RMSF for each residue represented by *C*_*α*_. The sharp peak around residue number 472 is due to the fluctuations of the unstructured loop. Due to the flexibility of the C-terminus tail domains, RMSF values are generally higher past the 625^th^ residue. Similar behavior is also seen for the first ∼ 14 residues which are more mobile than the residues immediately following them. (C) Side view of the N-terminal domains for apo (light blue) and +ssDNA (orange) show the rotation of domains in the +ssDNA system. Domain 1 is superimposed for both systems.

In apo equilibrium runs, the unstructured loop (Figure 1(A)) made contacts with the toroid interior around 140 ns mark. The loop interacted with the residues lining the cavity and the two interface alpha helices once inside the toroid. At the same time, a sliding motion of D1 against D3 occured, along with a large fluctuation of the C-terminus domains. The loop and D1-D3 motions combined were responsible for the jump in RMSD of the core N-terminal domains seen in Figure 2(A). Neither the D1-D3 sliding motion, or the abrupt large C-terminus fluctuation occured in the +ssDNA equilibration run. For +ssDNA, the unstructured loop quickly found itself in contact with the inner surface of the toroid preventing any large motions later on in the simulation. A comparison of apo system against +ssDNA at the end of equilibrium run is shown in Figure 2(C). The figure shows that in presence of ssDNA at the binding groove, the protein adopts a conformation where the D2 and D3 appear rotated forward. A similar rotation between apo and ssDNA bound topoisomerase structures was observed in a previous structural study [38].

The per-residue Root Mean Square Fluctuation (RMSF) plots in Figure 2(B) highlight some important motions in the protein structure. First, both curves demonstrate the high mobility of C-terminus domains in the equilibrium simulations. For both systems, the fluctuations for the C-terminal tail domains (grey, Figure 1) are higher past D4 residues (residue number 625). Likewise, the first ∼ 14 residues on the N-terminus are also highly mobile due to a lack of secondary structure. Another region that shows high fluctuations for both systems is the unstructured loop that exists in the vicinity of the toroid cavity. For apo, we see ∼ 4 fold increase in RMSF for residues comprising the loop vs residues in the vicinity of the loop. For +ssDNA, the fluctuations of the loop are lower, but still higher than the neighboring residues. The RMSF values are lower for the unstructured loop in the +ssDNA system as it finds a stable conformation quickly inside the cavity and does not fluctuate much more throughout the simulation time.

This unstructured loop is predicted to have a regulatory role during supercoil relaxation[14]. When the loop protrudes into the toroid cavity, it blocks the entrance into the toroid, and alters the surface electrostatics of the cavity rim. This loop is highly negatively charged as well. Figure 3(A-D) shows that in its absence, the cavity rim is electrostatically positive which is favorable for DNA binding. Its presence in the cavity however reduces the space available for DNA and also alters the charge of the interior. This was previously observed using *M. tuberculosis* Top1A where the presence or absence of the loop in the cavity altered the electrostatics in a similar manner [14]. This is significant in that both the loop and the ssDNA cannot simultaneously exist in the cavity. Multiple sequence alignment in Figure 3(E) shows that the loop is conserved for Mycobacterial type 1A topoisomerases but not other species. This presents an opportunity to target Mycobacteria specifically through the loop and cavity interactions.

**Figure 3:**
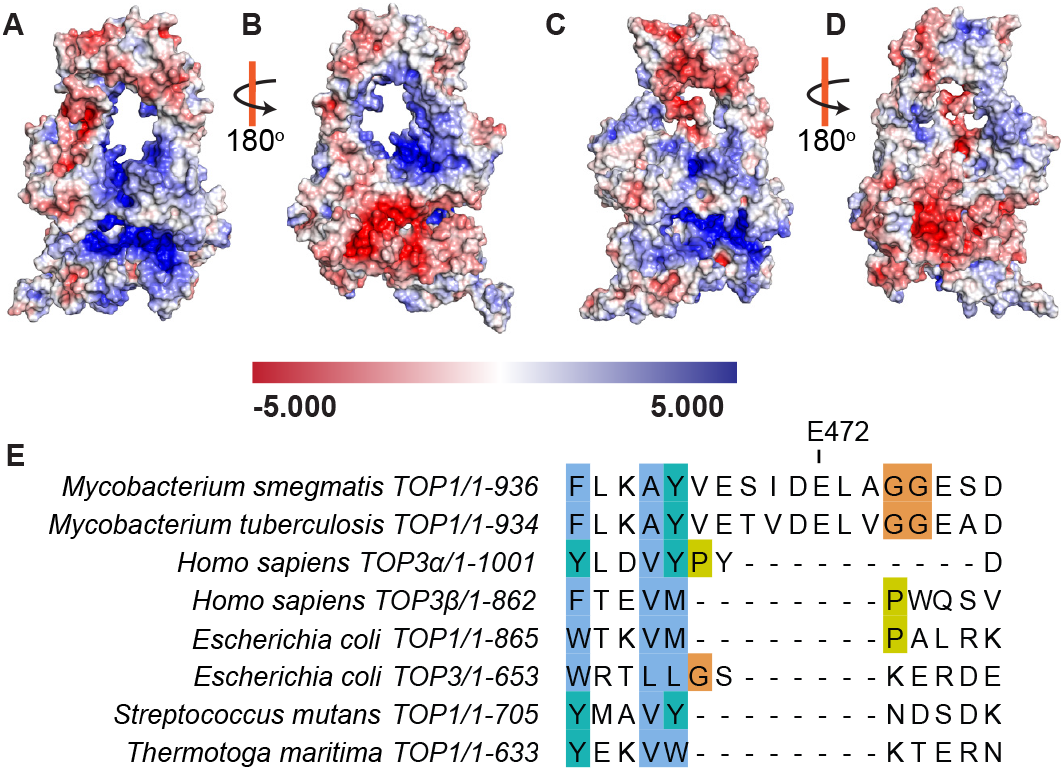
Electrostatic potential map for the Msm Top1A apo protein showing the effect of unstructured loop in the cavity and its conservation. (A-D) The apo system electrostatics showing the protein surface painted with colors representing the electrostatic potential on a scale of − 5*kT* / *e* (red) to 5*kT* / *e* (blue). Snapshot showing the front view (A) and back view (B) when the loop is not present in the toroid cavity. The blue regions depict areas of high DNA binding affinity. Cavity electrostatics is altered when the unstructured loop is present in the cavity as seen with the front view (C), and rear view (D). C-terminal tail domains are omitted for clarity. (E) Sequence alignment of type 1A topoisomerases from different species shown for the unstructured loop region. The loop insertion exists for *M. smegmatis* and *M. tuberculosis* but not the other type 1A topoisomerases included here. Residues colored using the Clustal coloring scheme where unconserved residues are colored white, hydrophobic residues are blue, Glycines are orange, Prolines are yellow, and aromatic residues are cyan.

### Stability of the gate-closed conformation

The majority of type 1A topoisomerases structures that are currently available are of the protein in the closed form either as apo or in complex with ssDNA strand(s). Previously single molecule experiments using Magnetic Tweezers found that the *E. coli* type 1A topoisomerases that were bound to ssDNA and formed covalent complexes were able to stay in a gate closed state against as much as 12 pN of opposing force[17]. This suggests that the protein highly favors a clamp closed state and is stable in this conformation. In our simulations, the core domains RMSD and RMSF show minimal fluctuations as well. To assess the stability of the gate closed state, we computed the total nonbonded energy (sum of pairwise electrostatic and van der Waals energy) of interaction between charged residues that formed contacts across the gate interface. The interaction energy for all identified pairs throughout equilibration runs was calculated using the VMD NAMDEnergy plugin. Snapshots of some of the interacting pairs are shown in Figure 4, while the time evolution is presented in Figure 5.

**Figure 4:**
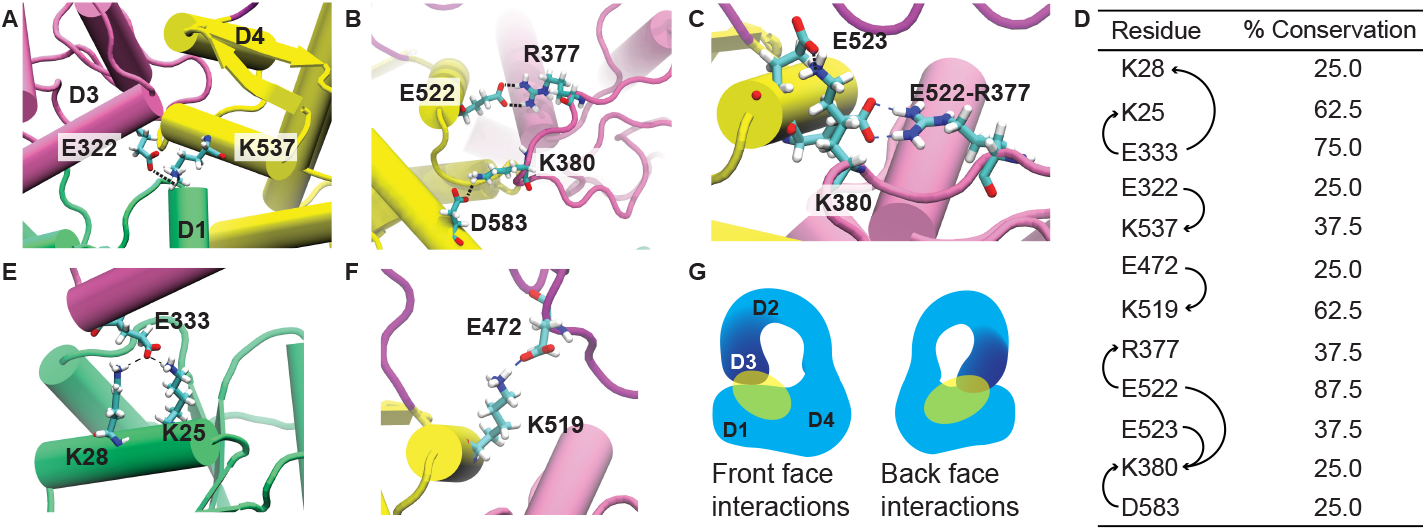
Nonbonded interactions between different residue pairs in front and back faces of the protein and their conservation. (A) In the front face of the protein, K537-E322 interaction is seen throughout the equilibrium runs for both systems. (B) In apo system, K380 forms strong contacts with D583, but not with E522. E522 interacts strongly with R377 instead. (C) In +ssDNA system, K380 interacts strongly with both E522 and E523. R377 also interacts with E522 at the same time. (E) In apo equilibrium runs, E333 forms contacts with K25 and K28 across the D1-D3 interface. This interaction is weak in the +ssDNA system. (F) The D2 unstructured loop can interact with charges lining the toroid interior. One such interaction is K519-E472. (G) Schematics of where these interactions occur on the protein surface. (D) Conservation of residues expressed in percentages when compared among the eight different type 1A topoisomerases as in Figure 3(E). Arrows depict residues that form contacts. Some residues are involved in contacts with multiple residues and have multiple arrows connected to them.

**Figure 5:**
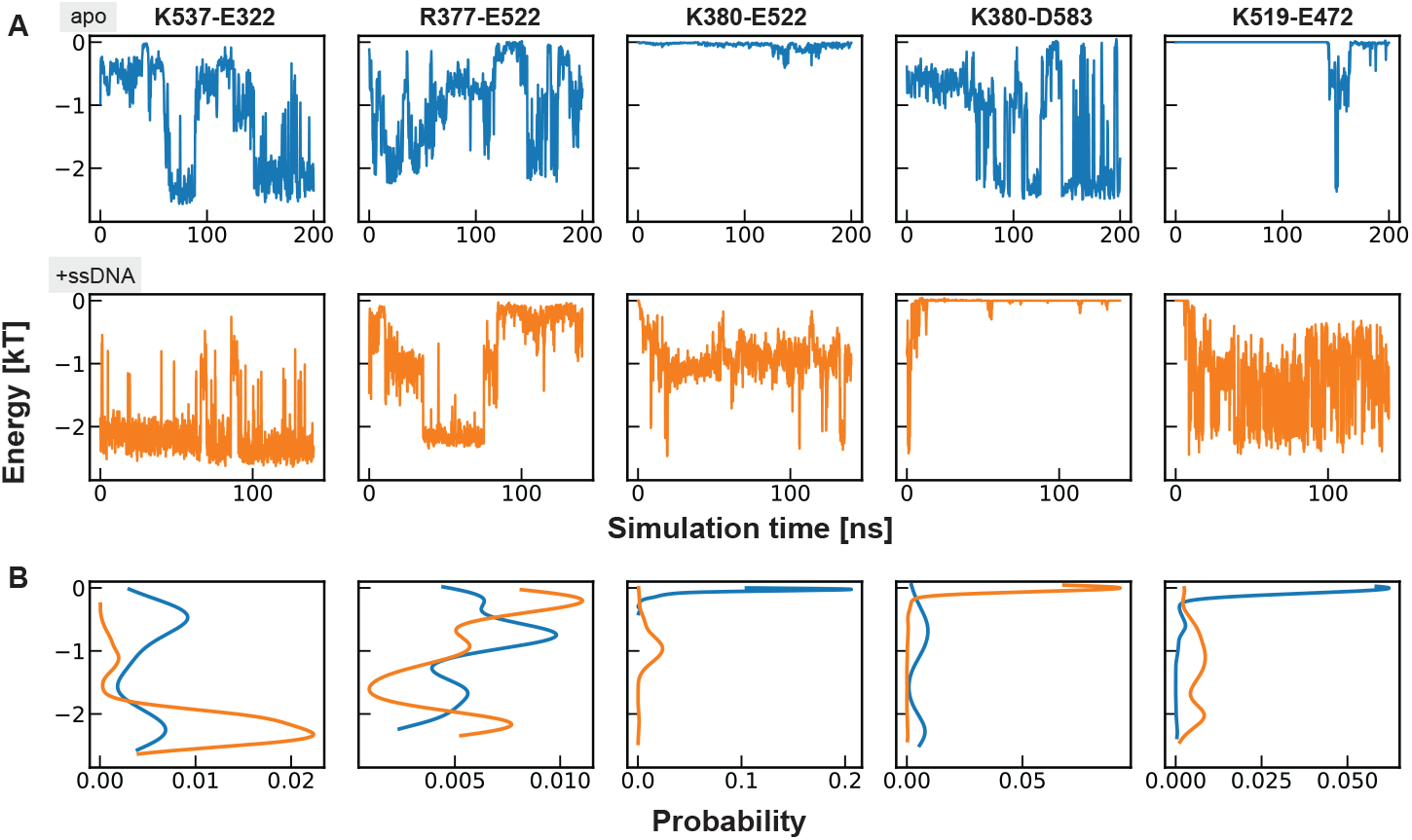
Nonbonded energy of interaction between different residue pairs. Top panels: apo, middle panels: +ssDNA, bottom panels: energy distributions for both apo and +ssDNA for each interacting pair. During the simulations, the strength of contacts varies dynamically. Comparison of the energies between different pairs show the relative strength of some interactions in comparison to others. Energies between other residue pairs mentioned in this work are plotted in Supplementary Figure 1 and 2.

From the time evolution and the distribution of nonbonded energies in Figure 5, several interesting features are apparent. First, we see that residue K537 (D4 *α*4) forms strong interactions with E322 (D3 *α*2) in both systems (see Supplementary figure 3 for the helices labeling). The proximity of the interacting residues is shown in Figure 4(A) while the strength of this interaction is plotted in Figure 5(A). This is a strong contact that likely contributes to the stability of the closed state. This contact exists in the front face of the protein.

Residues R377, K380, D583, E522 and E523 also form interactions that are important in stabilizing the gate closed structure. These residues form contacts on the back face of the protein. R377 and K380 from a D3 loop interact and form contacts with D4 residues E522, E523, and D583. In both systems, E522 (D4 *α*3) is able to form contacts with both R377 and K380. In apo, K380 forms strong contacts with D583 (D4 *α*6) while R377 interacts with E522. This is shown in Figure 4(B). In the +ssDNA system, the forward tilt of the D3 and D2 domains (Figure 2(C)) causes K380 to move away from D583 and instead interact strongly with E522 and E523. Across the D1-D3 interface, E333 forms strong contacts with K25 and K28 in the apo system, but not in the +ssDNA system. This is probably due to the interaction of ssDNA with charges in its vicinity. A small separation of D1 and D3 domain occurs at the interface (∼ 3*Å* in +ssDNA vs no gap in apo) due to lack of these contacts. In the toroid cavity, a strong interaction between K519 and E472 occurs around 150 ns in the apo system due to the D1-D3 slide along with the movement of the loop as mentioned previously. As shown in Figure 3(E), not all type 1A topoisomerases have this loop, so such an interaction could be unique to Mycobacteria.

We observed interaction of the same charged residue with multiple oppositely charged amino acids at several locations on the protein as well. As mentioned above, residue K380 interacts with multiple residues from D4 at the same time in the gate closed state. Likewise, E522 also interacts with both R377 and K380 in both systems. E522 is a conserved Glutamic acid among different type 1A topoisomerases (Figure 4(D)). Similar to findings in [14] (E527 for Mtb Top1A), E522 is likely important for Msm Top1A for activity and/or structure. In the interface between D1 and D3, E333 from D3 also forms interactions with two residues from D1 (Figure 4(E)). These redundant interactions between D3 and D1/D4 suggest a fail-safe mechanism to maintain the closed state structural integrity. In this manner, the protein might maintain its gate closed structure even accounting for issues such as mutations.

Sequence alignments were done to determine if the residues involved in these contacts are conserved across species. Sequence conservation expressed in percentages are displayed in Figure 4(D). All residues are conserved between topoisomerases of the two Mycobacteria. These residues all formed stabilizing interactions across the gate interface and contributed to the closed state stability for Msm Top1A. However, some residues that were not strictly conserved between species had opposite charged pair analogs such as in *Homo sapiens* Topoisomerase 3*α*(Hs Top3*α*). In Hs Top3*α* crystal structure (PDB ID: 4CHT), residues K345 and E543 are positioned close to one another in front of the protein similar to E322 and K537 for Msm Top1A. We speculate that K345-E543 is an interaction that contributes to the stability of the protein structure for Hs Top3*α* as well. Therefore, rather than strictly looking at the sequence conservation, it may be of greater importance to identify interaction pairs that similarly stabilize the closed state for different type 1A topoisomerases. This could be an interesting avenue for future studies.

### Umbrella sampling to sample the gate-open conformation

#### Umbrella sampling determines important domain motions during gate opening

To further probe the stability of the gate closed complex, we ran umbrella sampling to determine the free energy of gate opening. Umbrella sampling simulations were conducted from equilibrated structures as described in methods. The reaction coordinate (RC) was the distance between the center-of-mass (COM) of D3 and D1+D4 regions (Figure 6(A)). Increasing the reaction coordinate distance forces the protein to explore the gate-open state. Each system was steered from the closed to an open state by increasing the RC distance by 40*Å*. The gate is considered “open” when D1 residues detach from D3, and a gap arises between the D3 and D4 residues. For the activity of the protein, this gap must be large enough for a strand of DNA to pass through. Evidence has shown that type 1A topoisomerases open up during catalysis via a large conformational change and can fit even a dsDNA inside the cavity [11], [14].

**Figure 6:**
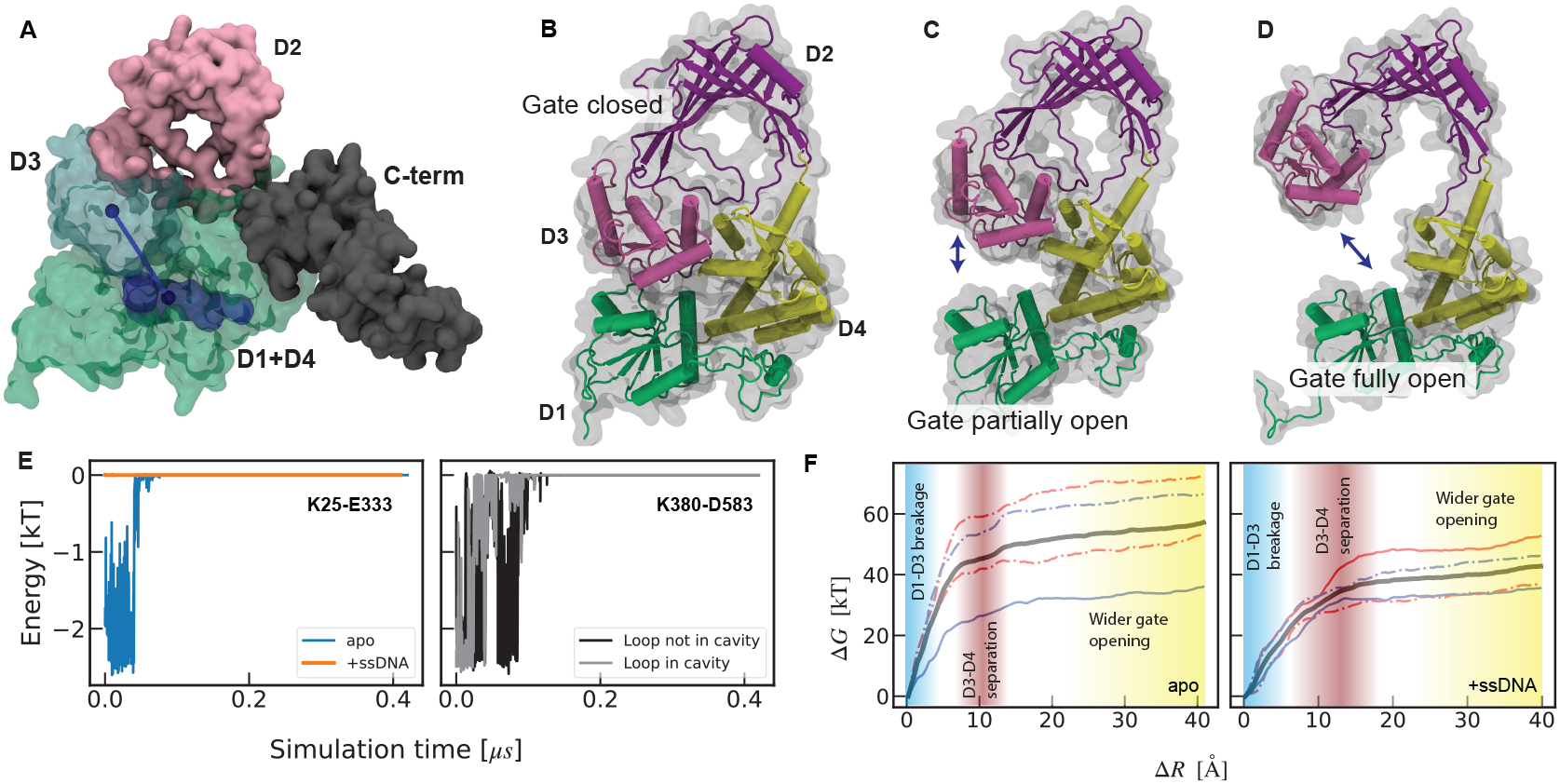
Umbrella sampling setup and results. (A) Reaction coordinate (solid blue line) for all US simulations is the distance between center of mass (COM) of the cyan region and the green region shown in solid blue spheres. The two regions contain D3 and D1+D4 residues respectively. ssDNA (blue) sits in a binding groove that spans D1 and D4. (B) Protein in a conformation where D1, D3, and D4 are in contact with one another. (C) During US runs, D1 detaches from D3 first. This is seen as a separation between the respective domains. (D) Finally, D3 detaches from D4 as further contacts are broken. Only N-terminus domains shown in all panels for clarity. (E) K25-E333 contact strength is strong in apo but negligible in +ssDNA system. K380-D583 contact stayed broken in presence of loop in the cavity. (F) Free energy curves generated using WHAM and the US outputs. Dashed-dotted lines in both plots represent systems with unstructured loop out of the cavity region(apo_1, or +ssDNA_1). ‘D1-D3 breakage’ means strictly the region where the gap between the two domains gets larger without the gate opening (panel C in this Figure). In the ‘D3-D4 separation’ region, the gate opens up. The final region ‘Wider gate opening’ refers to the space where the protein adopts a wider gate-open conformation with generally no interactions between any residues of D3 with D1 or D4. Thick black lines on both panels represent their respective average free energy of gate opening.

US simulations were conducted for apo and +ssDNA systems along with systems where the D2 unstructured loop was out of the cavity (apo_1 and +ssDNA_1). To ensure sufficient sampling, simulations were run with two different sampling times for both systems. The longer simulations are represented by blue curves in Figure 6(F) where the windows were sampled for 2.5*X* longer. Simulations with the unstructured loop out of the cavity are denoted by dash-dotted curves on the same figure.

For all US simulations, the pairwise nonbonded energies for contacts across the gate interface were computed as well. Contacts that stabilized the closed state broke as the gate opened. An example of interactions breaking during a US simulation is shown in Supplementary Figure 5. In some umbrella sampling simulations, occasionally there was formation of new interactions that were not present during equilibrium runs. Time evolution of two such interactions during an US run is shown in Supplementary Figure 6. Such interactions can alter the gate opening mechanisms by affecting the domain motions themselves. Still, all such interactions must break before the gate can fully open. Between different systems, time evolution of contacts show differences in their breakage behavior (Figure 6(E)). In apo, D1 and D3 are close to each other due to contacts between E333 and K25, resulting in a strong contact at the beginning of the simulation. The same interaction has negligible strength in the +ssDNA system which is readily broken. The presence of loop interactions in the cavity appeared to prevent reforming of strong K380-D583 contacts once broken in the apo system. This is one of the stronger contacts between D3 and D4. An earlier detachment of this contact would assist in gate opening.

Gate opening was seen to follow the same general motions in all US simulations. The first large motion was the movement of D1 down and away from D3 (Figure 6(C)). The separation between D1 and D3 was calculated as described in Methods. This distance between D1 and D3 residues before D3 detaches from D4 was found to be close in both set of systems ∼ (10 ± 4) *Å* for apo (and apo_1) and ∼ (12 ± 2) *Å* for +ssDNA (and +ssDNA_1). Without the movement of D1 away from D3 first, D2 could stretch sideways (as seen from front or back) (Supplementary Figure 4). This stretch could cause the gate to stay closed despite the interface helices being separated which opens the toroid cavity for DNA[11]. This D1-D3 separation is indicated on the free energy curves (Figure 6(F)) as first shaded region with blue gradient.

After D1 moves away from D3, the next motion is the detachment of D3 from D4 (Figure 6(D)). This separation of D3-D4 is indicated with the red gradient on both free energy curves 6(F). At this point, interactions across the gate interface involving residues such as K380, R377 and K537 break. This allows the gate to fully open. For systems with the unstructured loop in the cavity, this loop must also detach from any interactions across the gate interface. The loop usually sits over the cavity opening just above the two interface alpha helices (as shown in Figure 1(A) and Figure 6(B,C)), and therefore must move away from the interface before the opening is accessible. On average, separation of D3 and D4 takes place along the next 10*Å* of reaction coordinate distance. The separation of D3 beyond this point is referred to as “Wider gate opening” where the RC increases further representing a wide open DNA gate.

### US and WHAM generate free energies for the gate opening process

The Weighted Histogram Analysis Method(WHAM)[39] and US output were used to calculate the free energy of gate opening for all simulations (Figure 6(F)). In all cases, the free energy is funnel shaped with a deep well corresponding to the closed state and a plateau for the open state. This is consistent with the assumption that the open state is transient and the closed state is highly favored. For apo, longer sampling times did not always lower the barrier for gate opening (red vs blue dash-dotted curves). A comparison of the dash-dotted and solid blue curves show that despite having no unstructured loop interactions inside the cavity toroid during the gate opening, the free energy of this process for apo_1 was higher. For +ssDNA, neither the sampling time nor the presence of the unstructured loop affected the free energy curves significantly. A comparison of apo and +ssDNA simulations demonstrates that whenever the gate opens quickly via the separation of D3 from both D1 and D4 during the simulation run, the barrier heights are in general lower. This suggests that the barrier height is influenced by a combination of factors.

On average, the system with ssDNA present at the active site has lower barrier heights than the apo system. This may be because the ssDNA alters the electrostatics around the binding pocket and assists in the gate opening. Contacts K25-E333 and K28-E333 are both much weaker in the +ssDNA system than in apo (Supplementary figure 1, 2) suggesting that binding of the ssDNA highly affects the initial detachment of D1 from D3. Overall, this might assist in the easier detachment of the domains and result in an overall lower free energy barrier during the first crucial stages of gate opening.

The interactions of the D2 unstructured loop in the toroid did not significantly influence the free energy of gate opening either. As shown by the contrast between dash-dotted and solid curves in Figure 6(F), this difference alone was not enough to affect the energetics. The apo system results suggest that the loop might even assist in the gate opening by facilitating other motions during the process. However, the loop could be even more significant in regulating the movement of ssDNA to and from the toroid during the protein activity rather than the gate opening and closing processes. This remains to be confirmed.

### Principal Component Analysis

We saw in earlier sections that during US simulations for gate opening, different conformational changes and motions occur. Due to the strong interaction of atoms with their close environment, nearby atoms are forced to move in a concerted form. This means that the atomic motions are correlated. Principal Component Analysis (PCA) is a dimensionality reduction procedure that allows one to extract the functional motions of a molecule from such correlated motions [40]–[42]. We used PCA to determine major functional motions for all Msm Top1A umbrella sampling simulations.

We used the ensemble of protein conformations derived from each US simulation for PCA. For each simulation, the analysis found several components, but the top three components were retained as they were enough to describe most of the fluctuations of the system (>85%) (Supplementary Figure 7, Supplementary Note 1). Projection of the simulation trajectory on these components allows one to visualize the motions directly. The three components that were able to describe the majority of the protein’s motions were identified as (a) the detachment and downward motion of D1 away from D3, (b) the detachment of D3 from the D1+D4 interface and (c) swiveling of D3 out of the plane of the protein (Figure 7). Among different simulations, there was significant overlap of the components showing that similar motions were captured by the first three components in all simulations (Supplementary Table 1, Supplementary Note 2).

**Figure 7:**
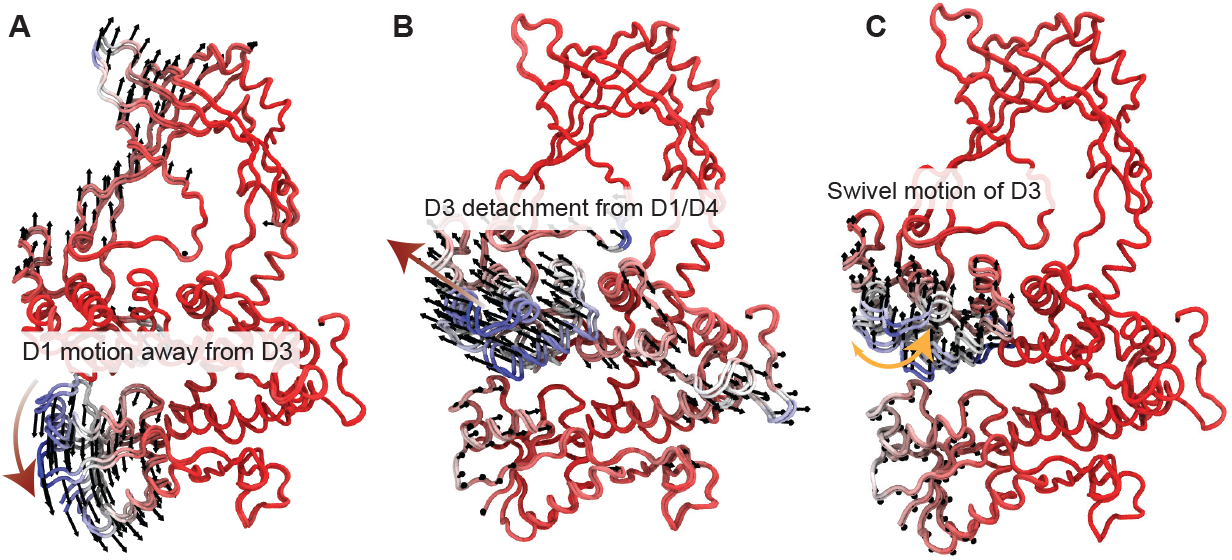
A few Principal Components describe most of the motions. (A), (B), and (C) show the three components that describe the majority of the system fluctuations as identified using Principal Component Analysis on a US run output. These three components from PCA resemble the major motions of the protein as it undergoes conformational changes from the gate-closed to a gate-open state.

We discussed the importance of D1-D3 detachment, as well as the D3 detachment from the D1 and D4 in the previous sections. We discuss the third identified motion - swivel of D3 and its importance in gate opening below.

### D3 swivel assists gate opening

One of the important functional motions identified from umbrella sampling simulations and principal component analysis is the swiveling of D3 during gate opening. Conformations seen during our umbrella sampling simulations (Figure 8(A,B)) are similar to the structure of a topoisomerase fragment previously observed in a swiveled state [43]. A recent cryo-EM gate-open structure of *Homo sapiens* Topoisomerase 3*β* (Hs Top3*β*) also shows substantial swivel of D2 and D3 together[16]. To investigate the extent of the swivel and its importance on gate opening during our simulations we computed 2D free energies. 2D free energies as a function of swiveling angles and reaction coordinate distance for both apo and +ssDNA systems was calculated using WHAM[30]. Figure 8(C) shows that the swiveling motion is most prominent after all interactions holding D3 close to D1 are broken. D3 is almost always observed swiveling outwards from the front face of the protein (see Figure 1(A) for the front face definition). We observe that D3 swiveling motion can occur either by just a rotation from the D2-D3 hinge region or along with a rotation of D2 and D3 together through the D2-D4 hinge. These hinge regions are comprised of loop structures and have been focused on several studies [14], [16], [38].

**Figure 8:**
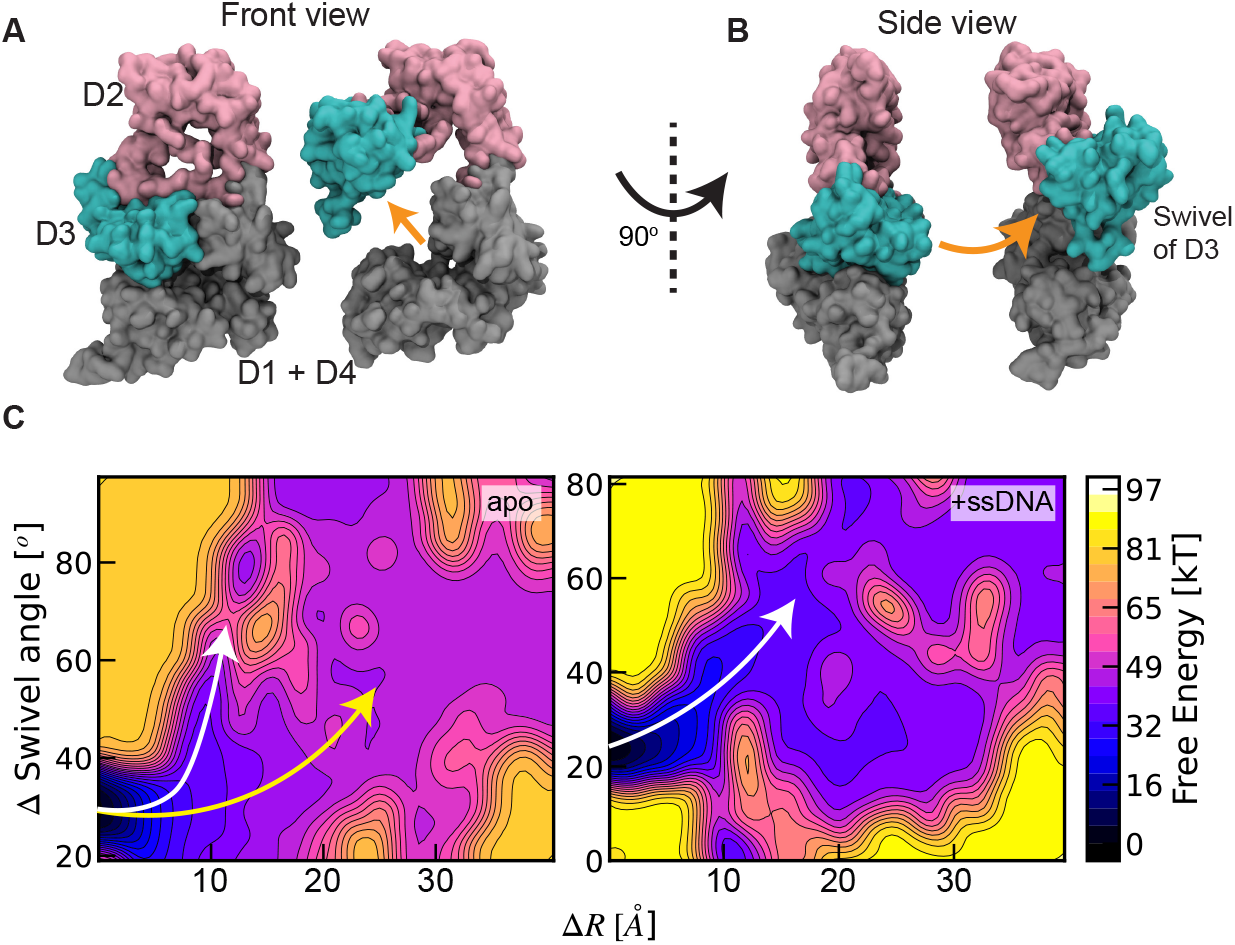
Swivel of D3 is important in the overall gate opening process. (A) Example snapshot of front view of a gate open structure. (B) Rotating (A) by 90^°^, one can see the large swivel of domain 3. Such swivel is observed in all gate opening simulations. (C) 2D free energy plots for apo (left) and +ssDNA (right) systems showing the energetic cost of changes in swiveling angle as a function of the reaction coordinate distance. Angle changes are normalized against the lowest starting swivel angle in both sets of simulations. Areas denoted by light yellow color are areas of very high free energy barrier not visited during the simulations. Angle changes are minimal during the first 5*Å* in both systems, but with a steep energy cost. The swiveling past 10 *Å* for both systems is not as energetically expensive. White arrow in both panels depict paths with lower energetic barrier to gate opening.

We note from Figure 8(C) that significant increases in swivel angles start around 5*Å* ≤ Δ*R* ≤ 8*Å* for +ssDNA. For apo, such increases start around Δ*R* ∼ 10*Å* with the exception of one simulation where an early increase in swivel angle was observed. This is the same simulation that resulted in the lower energetic barrier for gate opening (Figure 6(F), apo, solid blue curve) and comparable with the +ssDNA curves. Only this simulation proceeded along the path traced by the white line (Figure 8(C)). These Δ*R* locations roughly correlate with locations in the collective variable space where the gate begins to open up fully as seen in the free energy plots in Figure 6(F). Comparing Figure 6(F) and Figure 8(C), we see that the energetic cost of D3 swiveling motion is not as high as it is during the initial detachment of D1 and D3 and the separation of D3 from D4. Likewise, comparisons between the two Figures 6(F) and 8(C) also show that presence of ssDNA at the binding pocket facilitates earlier swiveling of the D3, lowering the free energy to gate opening. This suggests that the swiveling motion is important in gate opening, and such a motion is assisted by the binding of the ssDNA at the pocket.

This hinge flexibility and swiveling is also consistent with a proposed revised supercoil relaxation mechanism [12] and the recent structure [16]. In the first mechanism, it was proposed that D3 moves down towards D1 and D4 to capture the T-strand. This process necessitates that the hinges between D2 and both D3 and D4 be flexible. From Figure 8(C), we see that the hinges are quite flexible to allow such a motion. They allow for swiveling similar to observations from [43] and [16], in excess of 20^°^ previously seen in [38] but comparable to 60^°^ reported in[16]. As proposed in[14], our study demonstrates that the dissociation of D3 from D1 and D4 is an important step in gate opening. After detachment of the domains, a more pronounced swiveling out motion of D3 is seen where the residues move out of the plane of the protein.

## CONCLUSIONS

Type 1A topoisomerases undergo large conformational changes on short timescales. This requires dynamic experimental and computational methods to resolve the intricate details along their reaction cycle. The molecular dynamics studies presented here have revealed possible structural mechanisms of gate-opening - a major elusive state along the catalytic cycle.

Msm Top1A activity via strand-passage requires an opening between the D1, D3 and D4 interface domains. The interface is stabilized by electrostatic interactions between charged residues, all of which are conserved in Mycobacteria, and some across species (Figure 4(D)). During biased runs using umbrella sampling, these interactions are broken. This breaking of inter domain electrostatic interactions results in a gate-open structure that is consistent with several findings and predictions[2], [6], [14], [16], [17]. Our results indicate gate opening does not only involve the separation between domains, but also a swiveling of D3, congruent with other experimental findings [12], [16], [43].

Mutagenesis studies targeting the important residues and their interactions from this study might lead to further insights. Specifically, supercoil relaxation experiments and force dependent kinetics experiments would clarify the significance of these interactions as it pertains to protein activity. The approach described in this article could also be generalized to other type 1A topoisomerases – especially *Homo sapiens* type 1A topoisomerases. Structural studies into Hs topoisomerase 3*α* and 3*β* have garnered much interest in recent years and MD simulations could facilitate a similar study of structural transitions that occur during gate opening. These investigations could pave the way for further research into the general mechanisms and structural details of proteins belonging to the type 1A topoisomerase subfamily.

## AUTHOR CONTRIBUTIONS

D.S.: designed and performed research, analyzed data and wrote the manuscript. M.M.: designed research and wrote manuscript.

## ACKNOWLEDGMENTS

Parts of simulation work was performed using the high performance computing infrastructure provided by Research Support Services at the University of Missouri, Columbia MO. DOI: 10.32469/10355/97710. This work was supported by startup funds from University of Missouri Department of Physics and Astronomy to M.M.

## REFERENCES

[1] S. J. McKie, K. C. Neuman, and A. Maxwell, “DNA topoisomerases: Advances in understanding of cellular roles and multi-protein complexes via structure-function analysis,”

[2] A. J. Schoeffler and J. M. Berger, “DNA topoisomerases: Harnessing and constraining energy to govern chromosome topology,” Quarterly Reviews of Biophysics, vol. 41, no. 1, pp. 41–101, Feb. 2008. doi: 10.1017/S003358350800468X.

[3] B. Diaz, C. Mederos, K. Tan, and Y.-C. Tse-Dinh, “Microbial type IA topoisomerase c-terminal domain sequence motifs, distribution and combination,” International Journal of Molecular Sciences, vol. 23, no. 15, p. 8709, Aug. 5, 2022. doi: 10.3390/ijms23158709.

[4] C. D. Lima, J. C. Wang, and A. Mondragón, “Three-dimensional structure of the 67k n-terminal fragment of e. coli DNA topoisomerase i,” Nature, vol. 367, no. 6459, pp. 138–146, Jan. 1994. doi: 10.1038/367138a0.

[5] J. J. Champoux, “DNA topoisomerases: Structure, function, and mechanism,” Annual Review of Biochemistry, vol. 70, no. 1, pp. 369–413, Jun. 2001. doi: 10.1146/annurev.biochem.70.1.369.

[6] T. Dasgupta, S. Ferdous, and Y.-C. Tse-Dinh, “Mechanism of type IA topoisomerases,” Molecules, vol. 25, no. 20, p. 4769, Oct. 17, 2020. doi: 10.3390/molecules25204769.

[7] N. Cao, K. Tan, T. Annamalai, A. Joachimiak, and Y.-C. Tse-Dinh, “Investigating mycobacterial topoisomerase i mechanism from the analysis of metal and DNA substrate interactions at the active site,” Nucleic Acids Research, vol. 46, no. 14, pp. 7296–7308, Aug. 21, 2018. doi: 10.1093/nar/gky492.

[8] N. H. Dekker, V. V. Rybenkov, M. Duguet, et al., “The mechanism of type IA topoisomerases,” Proceedings of the National Academy of Sciences, vol. 99, no. 19, pp. 12 126–12 131, Sep. 17, 2002. doi: 10.1073/pnas.132378799.

[9] K. Terekhova, J. F. Marko, and A. Mondragón, “Studies of bacterial topoisomerases i and III at the single-molecule level,” Biochemical Society Transactions, vol. 41, no. 2, pp. 571–575, Apr. 1, 2013. doi: 10.1042/BST20120297.

[10] M. J. Szafran, T. Strick, A. Strzałka, J. Zakrzewska-Czerwińska, and D. Jakimowicz, “A highly processive topoisomerase i: Studies at the single-molecule level,” Nucleic Acids Research, vol. 42, no. 12, pp. 7935–7946, Jul. 8, 2014. doi: 10.1093/nar/gku494.

[11] Z. Li, A. Mondragón, and R. J. DiGate, “The mechanism of type IA topoisomerase-mediated DNA topological transformations,” Molecular Cell, vol. 7, no. 2, pp. 301–307, Feb. 2001. doi: 10.1016/S1097-2765(01)00178-2.

[12] K. H. Gunn, J. F. Marko, and A. Mondragón, “An orthogonal single-molecule experiment reveals multiple-attempt dynamics of type IA topoisomerases,” Nature Structural & Molecular Biology, vol. 24, no. 5, pp. 484–490, May 2017. doi: 10.1038/nsmb.3401.

[13] A. Changela, R. J. DiGate, and A. Mondragón, “Structural studies of e. coli topoisomerase III-DNA complexes reveal a novel type IA topoisomerase-DNA conformational intermediate,” Journal of Molecular Biology, vol. 368, no. 1, pp. 105–118, Apr. 2007. doi: 10.1016/j.jmb.2007.01.065.

[14] S. Ferdous, T. Dasgupta, T. Annamalai, K. Tan, and Y.-C. Tse-Dinh, “The interaction between transport-segment DNA and topoisomerase IA—crystal structure of MtbTOP1 in complex with both g- and t-segments,” Nucleic Acids Research, vol. 51, no. 1, pp. 349–364, Jan. 11, 2023. doi: 10.1093/nar/gkac1205.

[15] N. Cao, K. Tan, X. Zuo, T. Annamalai, and Y.-C. Tse-Dinh, “Mechanistic insights from structure of mycobacterium smegmatis topoisomerase i with ssDNA bound to both n- and c-terminal domains,” Nucleic Acids Research, vol. 48, no. 8, pp. 4448–4462, May 7, 2020. doi: 10.1093/nar/gkaa201.

[16] X. Yang, X. Chen, W. Yang, and Y. Pommier, “Structural insights into human topoisomerase 3 DNA and RNA catalysis and nucleic acid gate dynamics,” Nature Communications, vol. 16, no. 1, p. 834, Jan. 19, 2025, Publisher: Nature Publishing Group. doi: 10.1038/s41467-025-55959-y.

[17] M. Mills, Y.-C. Tse-Dinh, and K. C. Neuman, “Direct observation of topoisomerase IA gate dynamics,” Nature Structural & Molecular Biology, vol. 25, no. 12, pp. 1111–1118, Dec. 2018. doi: 10.1038/s41594-018-0158-x.

[18] J. A. M. Bakx, A. S. Biebricher, G. A. King, et al., “Duplex DNA and BLM regulate gate opening by the human TopoIII-RMI1-RMI2 complex,” Nature Communications, vol. 13, no. 1, p. 584, Dec. 2022. doi: 10.1038/s41467-022-28082-5.

[19] S. A. Hollingsworth and R. O. Dror, “Molecular dynamics simulation for all,” Neuron, vol. 99, no. 6, p. 1129, Sep. 9, 2018, Publisher: NIH Public Access. doi: 10.1016/j.neuron.2018.08.011.

[20] S. Ravishankar, A. Ambady, D. Awasthy, et al., “Genetic and chemical validation identifies mycobacterium tuberculosis topoisomerase i as an attractive anti-tubercular target,” Tuberculosis, vol. 95, no. 5, pp. 589–598, Sep. 2015. doi: 10.1016/j.tube.2015.05.004.

[21] V. Nagaraja, A. A. Godbole, S. R. Henderson, and A. Maxwell, “DNA topoisomerase i and DNA gyrase as targets for TB therapy,” Drug Discovery Today, vol. 22, no. 3, pp. 510–518, Mar. 2017. doi: 10.1016/j.drudis.2016.11.006.

[22] E. C. Meng, T. D. Goddard, E. F. Pettersen, et al., “UCSF ChimeraX: Tools for structure building and analysis,” Protein Science, vol. 32, no. 11, e4792, 2023, _eprint: https://onlinelibrary.wiley.com/doi/pdf/10.1002/pro.4792. doi: 10.1002/pro.4792.

[23] J. C. Phillips, D. J. Hardy, J. D. C. Maia, et al., “Scalable molecular dynamics on CPU and GPU architectures with NAMD,” The Journal of Chemical Physics, vol. 153, no. 4, p. 044 130, Jul. 28, 2020. doi: 10.1063/5.0014475.

[24] J. C. Phillips, R. Braun, W. Wang, et al., “Scalable molecular dynamics with NAMD,” Journal of Computational Chemistry, vol. 26, no. 16, pp. 1781–1802, 2005, _eprint: https://onlinelibrary.wiley.com/doi/pdf/10.1002/jcc.20289. doi: 10.1002/jcc.20289.

[25] J. Huang and A. D. MacKerell Jr, “CHARMM36 all-atom additive protein force field: Validation based on comparison to NMR data,” Journal of Computational Chemistry, vol. 34, no. 25, pp. 2135–2145, 2013, _eprint: https://onlinelibrary.wiley.com/doi/pdf/10.1002/jcc.23354. doi: 10.1002/jcc.23354.

[26] R. J. Gowers, M. Linke, J. Barnoud, et al., “MDAnalysis: A python package for the rapid analysis of molecular dynamics simulations,” Proceedings of the 15th Python in Science Conference, pp. 98–105, 2016, Conference Name: Proceedings of the 15th Python in Science Conference. doi: 10.25080/Majora-629e541a-00e.

[27] N. Michaud-Agrawal, E. J. Denning, T. B. Woolf, and O. Beckstein, “MDAnalysis: A toolkit for the analysis of molecular dynamics simulations,” Journal of Computational Chemistry, vol. 32, no. 10, pp. 2319–2327, 2011, _eprint: https://onlinelibrary.wiley.com/doi/pdf/10.1002/jcc.21787. doi: 10.1002/jcc.21787.

[28] G. M. Torrie and J. P. Valleau, “Nonphysical sampling distributions in monte carlo free-energy estimation: Umbrella sampling,” Journal of Computational Physics, vol. 23, no. 2, pp. 187–199, Feb. 1, 1977. doi: 10.1016/0021-9991(77)90121-8.

[29] B. Roux, “The calculation of the potential of mean force using computer simulations,” Computer Physics Communications, vol. 91, no. 1, pp. 275–282, Sep. 2, 1995. doi: 10.1016/0010-4655(95)00053-I.

[30] A. Grossfield, “An implementation of WHAM: The weighted histogram analysis method,” vol. 2.0.10,

[31] S. Zhang, J. M. Krieger, Y. Zhang, et al., “ProDy 2.0: Increased scale and scope after 10 years of protein dynamics modelling with python,” Bioinformatics, vol. 37, no. 20, pp. 3657–3659, Oct. 25, 2021. doi: 10.1093/bioinformatics/btab187.

[32] A. Bakan, L. M. Meireles, and I. Bahar, “ProDy: Protein dynamics inferred from theory and experiments,” Bioinformatics, vol. 27, no. 11, pp. 1575–1577, Jun. 1, 2011. doi: 10.1093/bioinformatics/btr168.

[33] A. Bakan, A. Dutta, W. Mao, et al., “Evol and ProDy for bridging protein sequence evolution and structural dynamics,” Bioinformatics, vol. 30, no. 18, pp. 2681–2683, Sep. 15, 2014. doi: 10.1093/bioinformatics/btu336.

[34] W. Humphrey, A. Dalke, and K. Schulten, “VMD: Visual molecular dynamics,” 1996. doi: 10.1016/0263-7855(96)00018-5.

[35] C. Camacho, G. M. Boratyn, V. Joukov, R. Vera Alvarez, and T. L. Madden, “ElasticBLAST: Accelerating sequence search via cloud computing,” BMC Bioinformatics, vol. 24, no. 1, p. 117, Mar. 26, 2023. doi: 10.1186/s12859-023-05245-9.

[36] A. M. Waterhouse, J. B. Procter, D. M. A. Martin, M. Clamp, and G. J. Barton, “Jalview version 2—a multiple sequence alignment editor and analysis workbench,” Bioinformatics, vol. 25, no. 9, pp. 1189–1191, May 1, 2009. doi: 10.1093/bioinformatics/btp033.

[37] J. D. Hunter, “Matplotlib: A 2d graphics environment,” Computing in Science & Engineering, vol. 9, no. 3, pp. 90–95, May 2007, Conference Name: Computing in Science & Engineering. doi: 10.1109/MCSE.2007.55.

[38] A. Changela, R. J. DiGate, and A. Mondragón, “Crystal structure of a complex of a type IA DNA topoisomerase with a single-stranded DNA molecule,” Nature, vol. 411, no. 6841, pp. 1077–1081, Jun. 2001. doi: 10.1038/35082615.

[39] A. Grossfield, WHAM: The weighted histogram analysis method, version 2.0.11.

[40] J. Palma and G. Pierdominici-Sottile, “On the uses of PCA to characterise molecular dynamics simulations of biological macromolecules: Basics and tips for an effective use,” ChemPhysChem, vol. 24, no. 2, e202200491, 2023, _eprint: https://onlinelibrary.wiley.com/doi/pdf/10.1002/cphc.202200491. doi: 10.1002/cphc.202200491.

[41] L. Skjaerven, A. Martinez, and N. Reuter, “Principal component and normal mode analysis of proteins; a quantitative comparison using the GroEL subunit,” Proteins: Structure, Function, and Bioinformatics, vol. 79, no. 1, pp. 232–243, 2011, _eprint: https://onlinelibrary.wiley.com/doi/pdf/10.1002/prot.22875. doi: 10.1002/prot.22875.

[42] D. R. Livesay, Ed., Protein Dynamics: Methods and Protocols, vol. 1084, Methods in Molecular Biology, Totowa, NJ: Humana Press, 2014. doi: 10.1007/978-1-62703-658-0.

[43] H. Feinberg and C. D. Lima, “Conformational changes in e. coli DNA topoisomerase i,” nature structural biology, vol. 6, no. 10, 1999.

