## Supplementary Materials for "Molecular dynamics simulations reveal DNA gate opening mechanisms for *M. smegmatis* topoisomerase 1A"

1. Apo equilibrium simulation, energy of interaction between residues

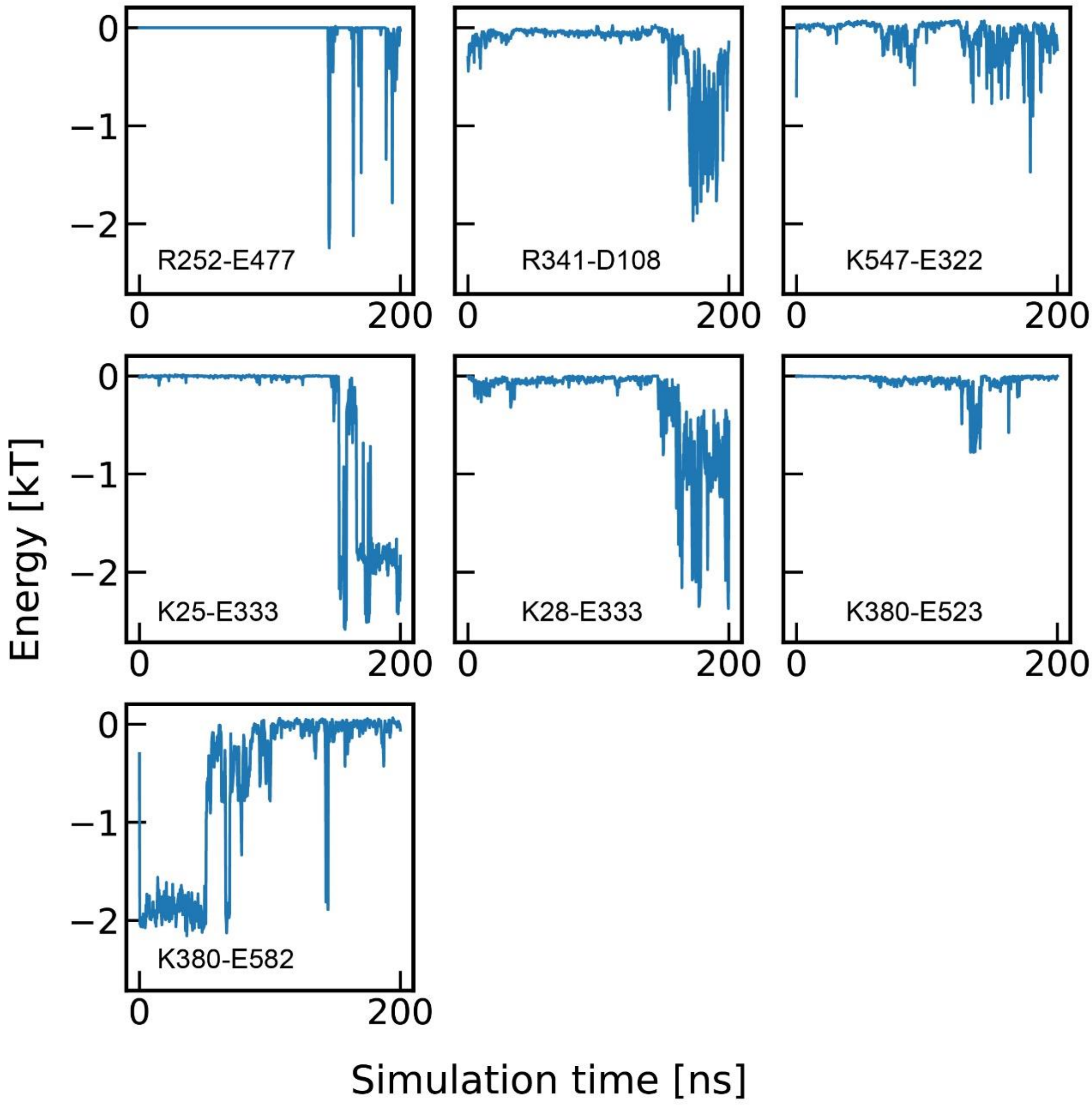


*Figure 1: Pairwise energy of interaction between residues in the apo equilibrium simulation.*

#### 2. +ssDNA equilibrium simulation, energy of interaction between residues

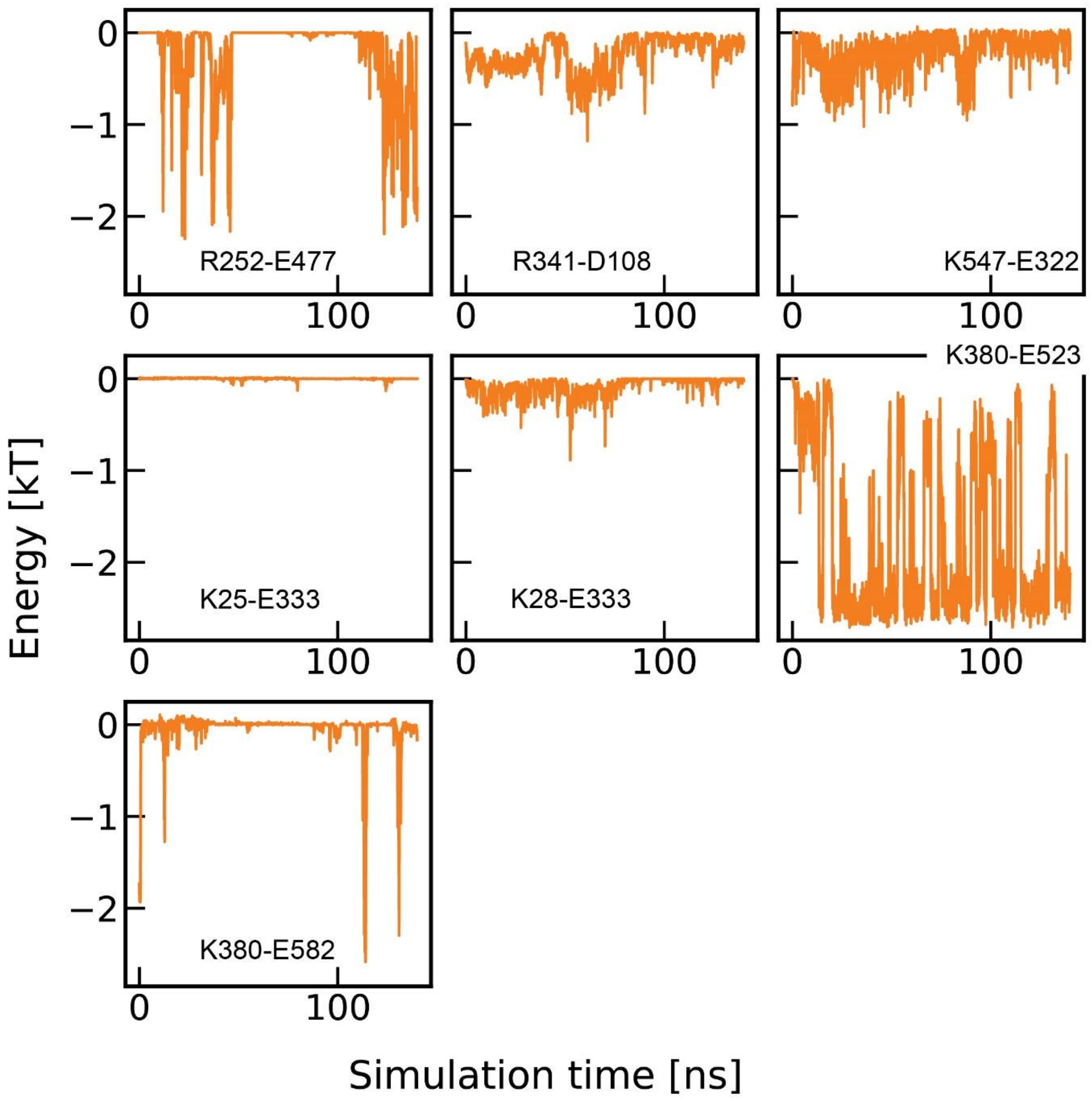


*Figure 2: Pairwise energy of interaction between residues in the +ssDNA equilibrium simulation.*

#### 3. Important helices

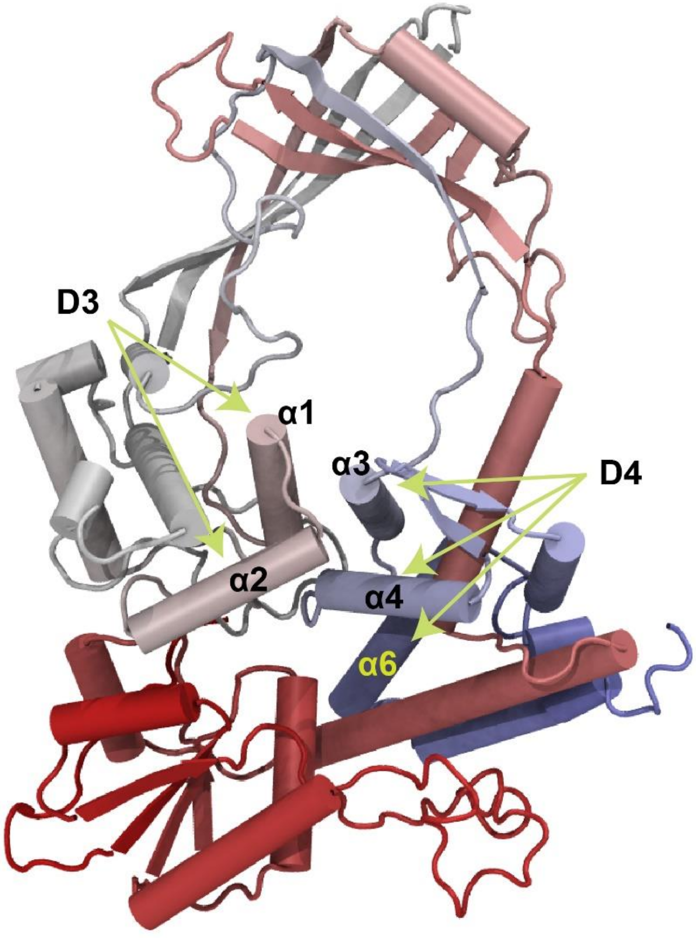


*Figure 3: Helices consisting of important residues mentioned in the text are labeled. Only N-terminus domains are shown for clarity.*

### 4. Distorted protein structures due to lack of D1-D3 separation

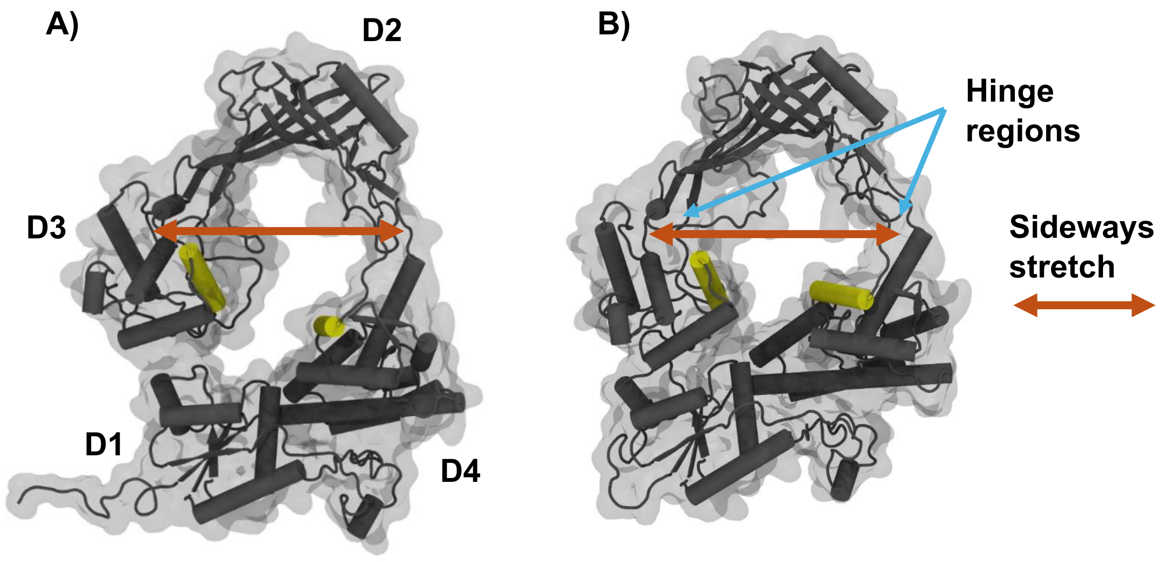


*Figure 4: Protein structures can get distorted during a biased simulation. The two yellow helices are the interface alpha helices. Their separation constitutes full gate opening [1]. However, if D1 and D3 fail to separate as seen here despite the separation of the interface helices, the gate is not fully open for supercoil relaxation. The protein may rather undergo a sideways stretch (dark orange arrows). D1-D3 separation is therefore crucial to prevent such a distortion of the protein structure.*

### 5. Breaking of interactions during an umbrella sampling simulation

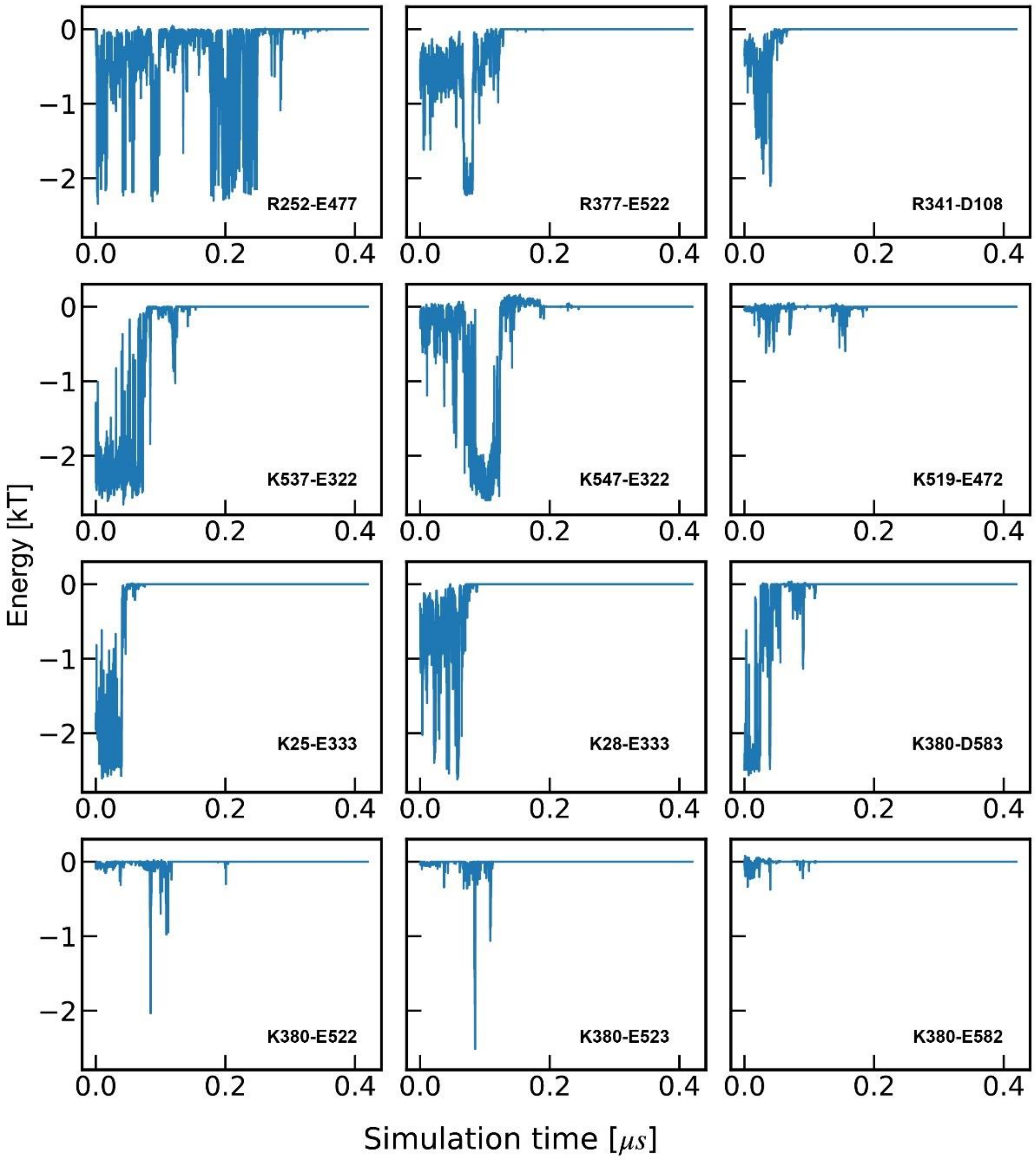


*Figure 5: Pairwise interaction energies calculated for an umbrella sampling simulation. Energies are calculated between the same residues identified during the equilibrium runs.*

### 6. Transient interactions form occasionally during an umbrella sampling simulation

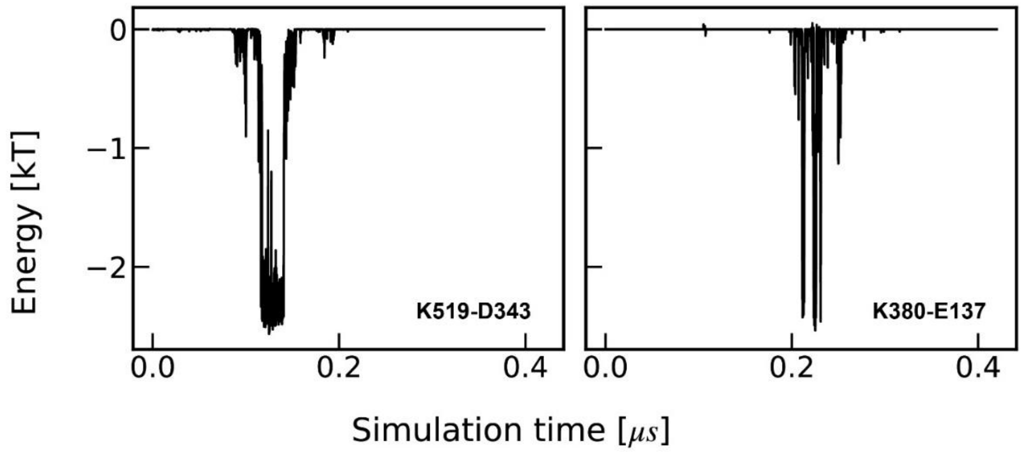


*Figure 6: Transient interactions that form during an umbrella sampling simulation. Such interactions can affect the gate opening by making newer connections between domains that were separated, making domain separation difficult, hindering or assisting D3 swivel and so on.*

### 7. PCA Eigenvalue Spectrum

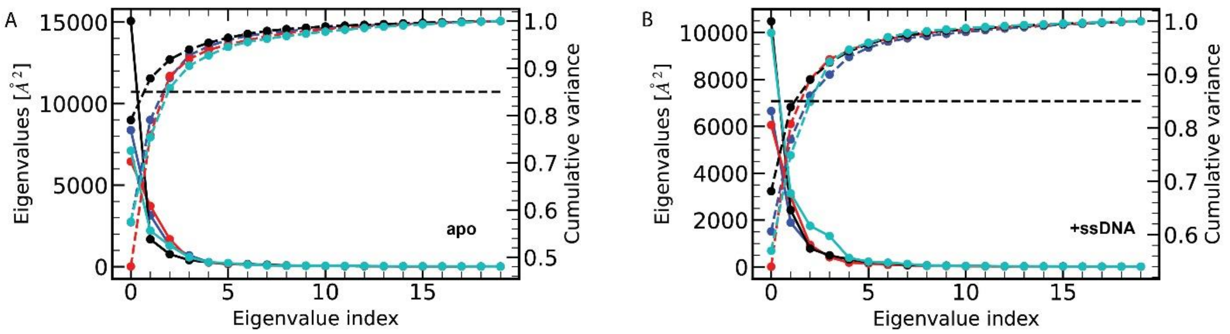


*Figure 7: (A) Eigenvalues calculated from 4 umbrella sampling simulations for apo (left y-axis) and their contributions to the cumulative variance (right y-axis). Twenty components are plotted. (B) Same as (A), but for +ssDNA system. Same colors represent the same simulation. Solid lines represent the eigenvalues, while dashed lines represent the cumulative variances. Black dashed line in both panels indicate the 85% cumulative variance mark.*

#### Supplementary Note 1:

The fluctuations of the systems are described by the eigenvalues along each eigenvectors (or components) generated from the analysis. The sharp initial decrease of the eigenvalues shows that the atomic motions are highly correlated. In all runs, the first three uncorrelated principal components represent > 85% of the total squared fluctuations. This indicates that the first few principal components can explain most of the total observed fluctuations in these simulations.

### 1. PCA RMSIP table for all umbrella sampling simulations

| | apo_1 | apo_1 | apo | apo_1 | +ssDNA_1 | +ssDNA | +ssDNA | +ssDNA_1 |
|---|---|---|---|---|---|---|---|---|
| **apo_1** | 1.00 | 0.90 | 0.79 | 0.79 | 0.47 | 0.48 | 0.50 | 0.43 |
| **apo_1** | | 1.00 | 0.72 | 0.81 | 0.57 | 0.57 | 0.59 | 0.53 |
| **apo** | | | 1.00 | 0.88 | 0.59 | 0.58 | 0.62 | 0.58 |
| **apo_1** | | | | 1.00 | 0.61 | 0.61 | 0.63 | 0.58 |
| **+ssDNA_1** | | | | | 1.00 | 0.84 | 0.85 | 0.72 |
| **+ssDNA** | | | | | | 1.00 | 0.90 | 0.70 |
| **+ssDNA** | | | | | | | 1.00 | 0.69 |
| **+ssDNA_1** | | | | | | | | 1.00 |

*Table 1: RMSIP values calculated between the subspace spanned by the first three components in all eight US simulations. apo_1 and +ssDNA_1 do not contain unstructured loop interactions in the toroid cavity. Black and orange colored text indicate RMSIP values between simulations from apo and +ssDNA system respectively. Blue colored text indicates the cross-simulation RMSIP values.*

#### Supplementary Note 2:

To quantify the degree of overlap between the first three components from all eight umbrella sampling simulations, we calculated the Root Mean Square Inner Product (RMSIP) between those subspaces. RMSIP values were calculated using ProDy libraries [2], [3] where a value of RMSIP = 0 means that vectors of two essential spaces are orthogonal to each other, and a RMSIP = 1 means that two essential spaces occupy the same subspace. For the three components that captured the most important fluctuations in all apo (and apo_1) runs, $RMSIP_{avg}$ = 0.81±0.07, and for +ssDNA (and +ssDNA_1) runs, $RMSIP_{avg}$ = 0.78±0.09. These values indicate that there is a high degree of overlap among the three modes identified through PCA calculations in each system and simulation. The RMSIP for cross terms between the different systems is slightly lower at 0.56±0.06 indicating some overlap. This is probably due to the presence of ssDNA at the binding site that influences the motions during gate opening for +ssDNA runs, but not for apo. Errors are standard deviations.

### References:


[1] Z. Li, A. Mondragón, and R. J. DiGate, “The Mechanism of Type IA Topoisomerase-Mediated DNA Topological Transformations,” *Mol. Cell*, vol. 7, no. 2, pp. 301–307, Feb. 2001, doi: 10.1016/S1097-2765(01)00178-2.

[2] A. Bakan, L. M. Meireles, and I. Bahar, “ProDy: Protein Dynamics Inferred from Theory and Experiments,” *Bioinformatics*, vol. 27, no. 11, pp. 1575–1577, Jun. 2011, doi: 10.1093/bioinformatics/btr168.

[3] A. Amadei, M. A. Ceruso, and A. Di Nola, “On the convergence of the conformational coordinates basis set obtained by the essential dynamics analysis of proteins’ molecular dynamics simulations,” *Proteins Struct. Funct. Bioinforma.*, vol. 36, no. 4, pp. 419–424, 1999, doi: 10.1002/(SICI)1097-0134(19990901)36:4<419::AID-PROT5>3.0.CO;2-U.
